# *In Situ* Microwave Fixation to Define the Terminal Rodent Brain Metabolome

**DOI:** 10.1101/2022.08.16.504166

**Authors:** Jelena A. Juras, Madison B. Webb, Lyndsay E.A. Young, Kia H. Markussen, Tara R. Hawkinson, Michael D. Buoncristiani, Kayli E. Bolton, Peyton T. Coburn, Meredith I. Williams, Lisa PY. Sun, William C. Sanders, Ronald C. Bruntz, Lindsey R. Conroy, Chi Wang, Matthew S. Gentry, Bret N. Smith, Ramon C. Sun

## Abstract

The brain metabolome directly connects to brain physiology and neuronal function. Brain glucose metabolism is highly heterogeneous among brain regions and continues postmortem. Therefore, challenges remain to capture an accurate snapshot of the physiological brain metabolome in healthy and diseased rodent models. To overcome this barrier, we employ a high-power focused microwave for the simultaneous euthanasia and fixation of mouse brain tissue to preserve metabolite pools prior to surgical removal and dissection of brain regions. We demonstrate exhaustion of glycogen and glucose and increase in lactate production during conventional rapid brain resection prior to preservation by liquid nitrogen that is not observed with microwave fixation. Next, microwave fixation was employed to define the impact of brain glucose metabolism in the mouse model of streptozotocin-induced type 1 diabetes. Using both total pool and isotope tracing analyses, we identified global glucose hypometabolism in multiple regions of the mouse brain, evidenced by reduced ^13^C enrichment into glycogen, glycolysis, and the TCA cycle. Reduced glucose metabolism correlated with a marked decrease in GLUT2 expression and several metabolic enzymes in unique brain regions. In conclusion, our study supports the incorporation of microwave fixation to study terminal brain metabolism in rodent models.

## INTRODUCTION

Brain metabolism is the biochemical basis of memory, cognition, and central nervous system (CNS)/peripheral neuronal signaling. The brain consumes approximately 20% of circulating glucose^1^ and although debated, neurons are proposed to be the major glucose-consuming cells of the brain^2,3^. Glucose metabolism of the brain can be subdivided into catabolic or anabolic processes. Catabolic processes involve the breakdown of glucose for energy production via glycolysis and the tricarboxylic acid cycle (TCA)^4^. Neurotransmitters such as GABA, glutamate, and aspartate, can be *de novo* synthesized through glucose catabolism^5^. In contrast, anabolic glucose metabolism utilizes either glucose or glucose derivatives for the synthesis of lipids^6^, complex carbohydrates such as glycans^7^, chondroitin^8^ and heparin sulfate biopolymers^9^, all of which are critical biomass and structural components of the brain. Given that brain glucose participates in a myriad of metabolic processes from bioenergetics to neurotransmitter and biomass production, it is difficult to decipher which downstream processes are perturbed from reduced 2-deoxyglucose uptake and are unique to disease pathogenesis. It is hypothesized that a deeper understanding of brain glucose metabolism will yield critical information for future treatment of multiple disorders such as diabetes, dementia, and epilepsy.

Glucose hypometabolism is characterized by reduced uptake of 2-fluorodeoxyglucose (FDG), imaged by Positron Emission Tomography (PET), and is a common pathological feature shared among multiple neurological disorders such as Alzheimer’s disease (AD)^10^, Parkinson’s disease^11^, temporal lobe epilepsy^12^, inherited pediatric neurodegenerative diseases^13^, and peripheral diseases such as type 1 diabetes^14^. Recently, the glycolysis-lactate axis has been highlighted as a key pathway perturbed in the disease-associated microglia during AD progression^15,16^. Further, we found that protein hyperglycosylation, an anabolic product of glucose metabolism, is a hallmark of AD in both human and mouse specimens^17^. Collectively, these studies highlight the potential of alternative metabolic fates of glucose in neurodegenerative diseases. Conversely, orphan neurological disorders such as Glut1-deficiency^13^, PDH-deficiency^18^, and Lafora disease all present with reduced glucose uptake on FDG-PET^19^, reduced oxidative phosphorylation and protein glycosylation, and patients are prone to epileptic seizures and accelerated neurodegeneration^18,20^. Therefore, clinical diagnosis of glucose hypometabolism and the downstream glucose metabolic alterations differ based on underlying disease pathology, highlighting the importance of accurately capturing physiological metabolite phenotypes in human and experimental models.

Currently, many techniques to accurately capture physiological brain metabolism in rodents *in vivo* rely on nuclear magnetic resonance (NMR)^21–23^, wherein mice are anesthetized, and live animals are used to study metabolic parameters. Metabolites captured by *in vivo* NMR are limited due to technical limitations and anesthesia could directly impacts brain metabolism^24,25^. Thus, large-scale metabolomics studies of the brain require rapid dissection of brain regions post-mortem followed by cryo-preservation and mass spectrometry detection of metabolites^20,26,27^. Post-mortem surgical removal of the brain ranges from 90 seconds to several minutes depending on the individual performing the resection. It is well documented that metabolism changes post-mortem^28,29^, and the rates of metabolic flux through glycolysis and the TCA cycle occurs in seconds^30–32^. Furthermore, artificial tissue hypoxia has been reported in mice following cervical dislocation^33^, decapitation^34^, and CO2 euthanasia^35^. Brain tissue collection through conventional routes results in glycogen degradation and changes in global protein phosphorylation status^36,37^. A deeper understanding of neuronal function hinges on expanded metabolic pathway coverage, creating a critical need to establish an accurate baseline brain metabolome that excludes residual non-physiological metabolism that occurs post-mortem brain resection.

Microwave fixation of mammalian tissues utilizes water molecule vibration to produce heat^38^, resulting in heat-inactivated protein denaturation, and in principle, stopping all metabolic processes^39^. Indeed, microwave fixation is frequently used in pathology laboratories to preserve mammalian tissues for histopathological analyses^40^. For direct *in situ* tissue fixation of whole animals, a focused microwave has been developed for the rapid inactivation of brain proteins (<0.6 seconds) and preserves global protein phosphorylation through enzyme inactivation^37^. Therefore, we hypothesized that enzymatic inactivation through microwave fixation would preserve the brain metabolome as well. Herein, we compared *in situ* microwave fixation to rapid dissection and cryo-preservation of wild-type C57/B6 brain tissues. We observed a drastic increase in free glucose utilization and glycogen degradation for lactate production during tissue removal and dissection of the brain using traditional cryo-preservation tissue harvesting. Further, we interrogated the impact of type 1 diabetes mellitus (T1DM) on brain metabolism using the streptozotocin (STZ)-induced hyperglycemia mouse model^41,42^, and observed profound glucose hypometabolism in the brain of T1DM animals using focused microwave.

## RESULTS

### *In situ* microwave fixation to define brain metabolism

Cellular metabolism continues post-mortem in rodents with signs of hypoxia driven metabolic reprogramming^29^. These metabolic changes are especially true for the brain, and capturing an accurate snapshot of the brain metabolome remains a critical challenge in the field. In this study, we utilized the focused microwave specifically designed for the rapid fixation of rodent brain tissue to prevent postmortem metabolic changes during brain resection. To test the application of focused microwave, we designed a two-arm study, where arm one is decapitation, followed by rapid dissection of the brain and crypreservation (Fig. 1). Arm two followed the similar workflow as arm 1, except brain tissue was fixed *in situ* using focused microwave prior to brain dissection (Fig. 1). The difference between the timing of metabolite fixation is 90 seconds (decapitation to cryo-preservation) and 0.6 seconds reported by the manufacture of focused microwave (Fig. 1A). Brain and other tissues from both cohorts of animals were pulverized while in liquid nitrogen by a cryomill and prepped for pooled metabolomics analysis by gas chromatography mass spectrometry (GCMS) for both polar metabolites and biomass^43–45^. For this study, we focused on the neocortex (CTX), hippocampus (HIPP), and the hindbrain dorsal vagal complex (DVC) which contains glucose sensing neurons that undergo functional neuroplasticity after prolonged hyperglycemia^46^. GCMS metabolomics analyses between focused microwave and cryo-preservation of brain regions displayed clear separation by Partial Least Squares-Discriminant Analysis (PLS-DA) (Fig. 1B), and metabolic pathway enrichment analysis suggesting changes in glycolysis, gluconeogenesis, TCA cycle, and amino acid metabolism that is shared between focused microwave and cryo-preservation among all three brain regions (Fig 1C, Table S1). Targeted metabolite analysis of the central carbon pathway revealed a drastic decrease in glycogen, glucose, citrate, and a profound increase in lactate, malate, and fumarate in the cryo-preservation arm (Fig 1D). Since the microwave beam of the focused microwave fixation does not have peripheral organs in its path, we included GCMS analysis of the lung, liver, and muscle as controls. We did not observe major changes between focused microwave and cryo-preservation methods with the exception of glucose and glucose-6-phosphate in liver and the muscle (Figure S1). Further, focused microwave shows improved variance within groups showing reduced coefficient of variance (CoV) among multiple metabolites (Fig. 1E). Finally, based on the current focused microwave data set (neocortex), n=5 / group would provide 89% power to detect those differences with 5% false discovery rate based on two-sample t-tests (Fig. 1F, Table S2).

**Figure 1.**
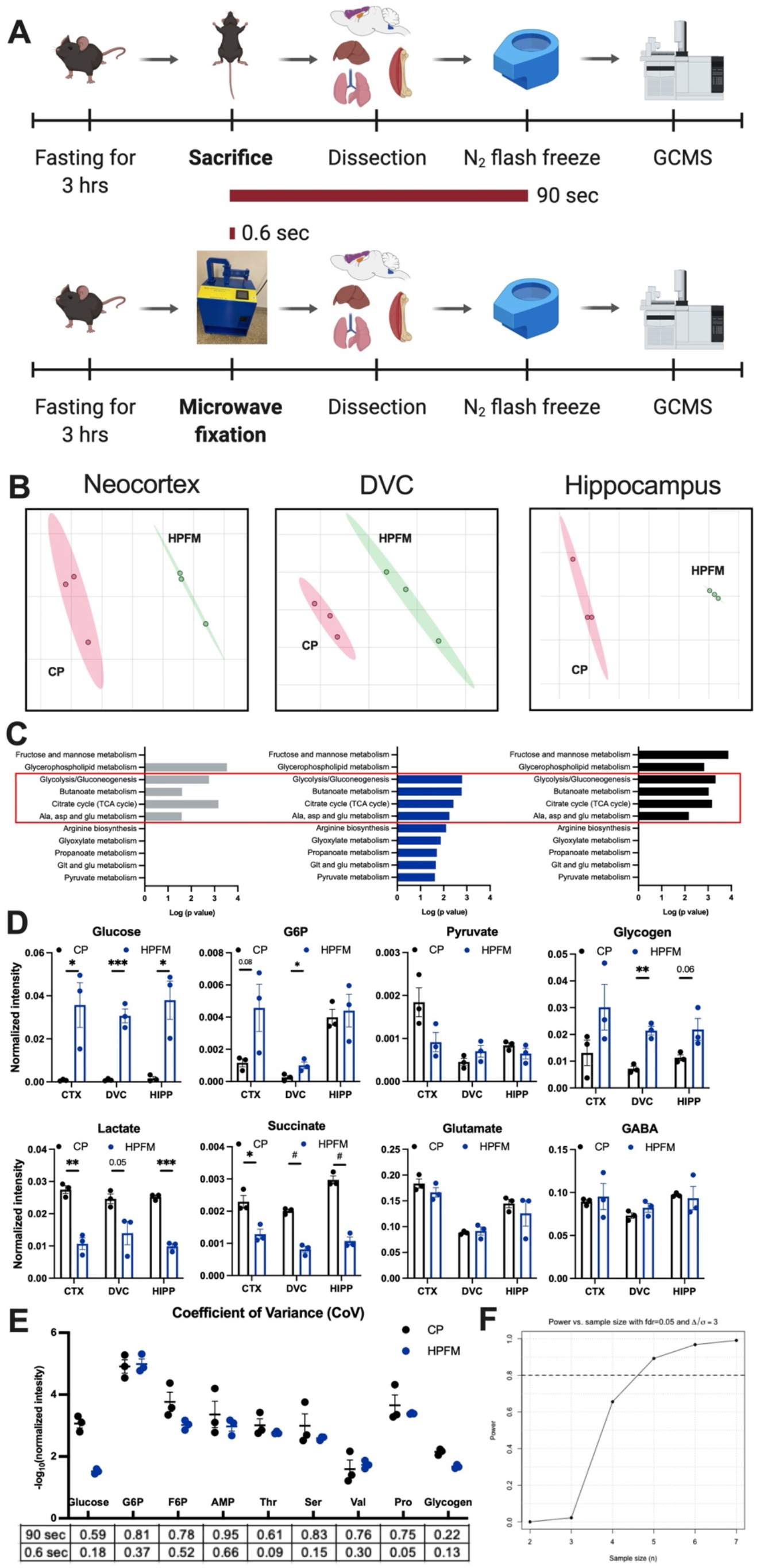
*In situ* high-power focused microwave fixation (HPFM) to study the brain metabolome. (**A**) Schematic of the experimental design. 6-week-old mice fasted for 3 hours, euthanized by decapitation (*top*) or microwave fixation (*bottom*), brain regions were then dissected, flash-frozen, and analyzed by GCMS. (**B**) Partial Least Squares-Discriminant Analysis (PLS-DA) of neocortex (CTX), dorsal vagal complex (DVC), and hippocampus (HIPP) metabolites showing distinct separation between mice euthanized by cryopreservation (CP) and HPFM. (**C**) Pathway analysis comparing HPFM and CP (performed by Metaboanalyst) of CTX, DVC, and HIPP. (**D**) Representatives metabolite levels from glycogen metabolism, glycolysis, and TCA cycle between CP and HPFM. Values are presented as mean ±SEM (n=3 biological replicates), *p<0.05, **p<0.01, ***p<0.001, #p<0.0001, analyzed by student t-test. (**E**) Coefficient of variance (CoV) analysis of representative metabolites in **B**. (**F**) Power analysis indicating the necessary sample size to detect 89% power at 0.05 false discovery rate. G6P: glucose 6-phosphate, F6P: fructose 6-phospahte, AMP: adenosine phosphate, Thr: Threonine, Ser: Serine, Val: Valine, Pro: Proline.

### *In situ* microwave fixation is compatible with stable isotope tracing

Stable isotopic tracing is invaluable at delineating substrate metabolism and interrogating enzymatic activities^47,48^. To test whether focused microwave is compatible with stable isotope tracing, we performed oral gavage delivery of ^13^C-glucose using a method previously described^44^ and performed the focused microwave and cryo-preservation, two-arm study in parallel to interrogate isotopic enrichment of central carbon metabolites between the two different brain fixative strategies (Fig 2A). We identified significantly increased enrichment of ^13^C in glycogen and glycolytic metabolites, including glucose, DHAP, and PEP in the neocortex following focused microwave (Fig 2B-C, Table S3); however, lactate, citrate, fumarate, malate, and amino acids such as glutamate, glutamine, serine, and GABA remain unchanged between focused microwave and cryo-preservation (Fig. 2C). It is likely that even though total pooled metabolites were changed during cryo-preservation, both ^13^C-enriched and ^12^C-TCA cycle and amino acid metabolites were either consumed or synthesized at the same rate during cryo-preservation. Similarly, we did not observe major changes in ^13^C-enrichment between focused microwave and cryo-preservation in the lung, liver, and SKM (Fig. S2, Table S4). Collectively these data suggest: 1) utilization of glycogen and glucose to supply glycolysis during the 90 seconds of tissue resection (Fig. 2D), and 2) stable isotope tracing could be a better strategy to define brain metabolism when focused microwave is not available.

**Figure 2.**
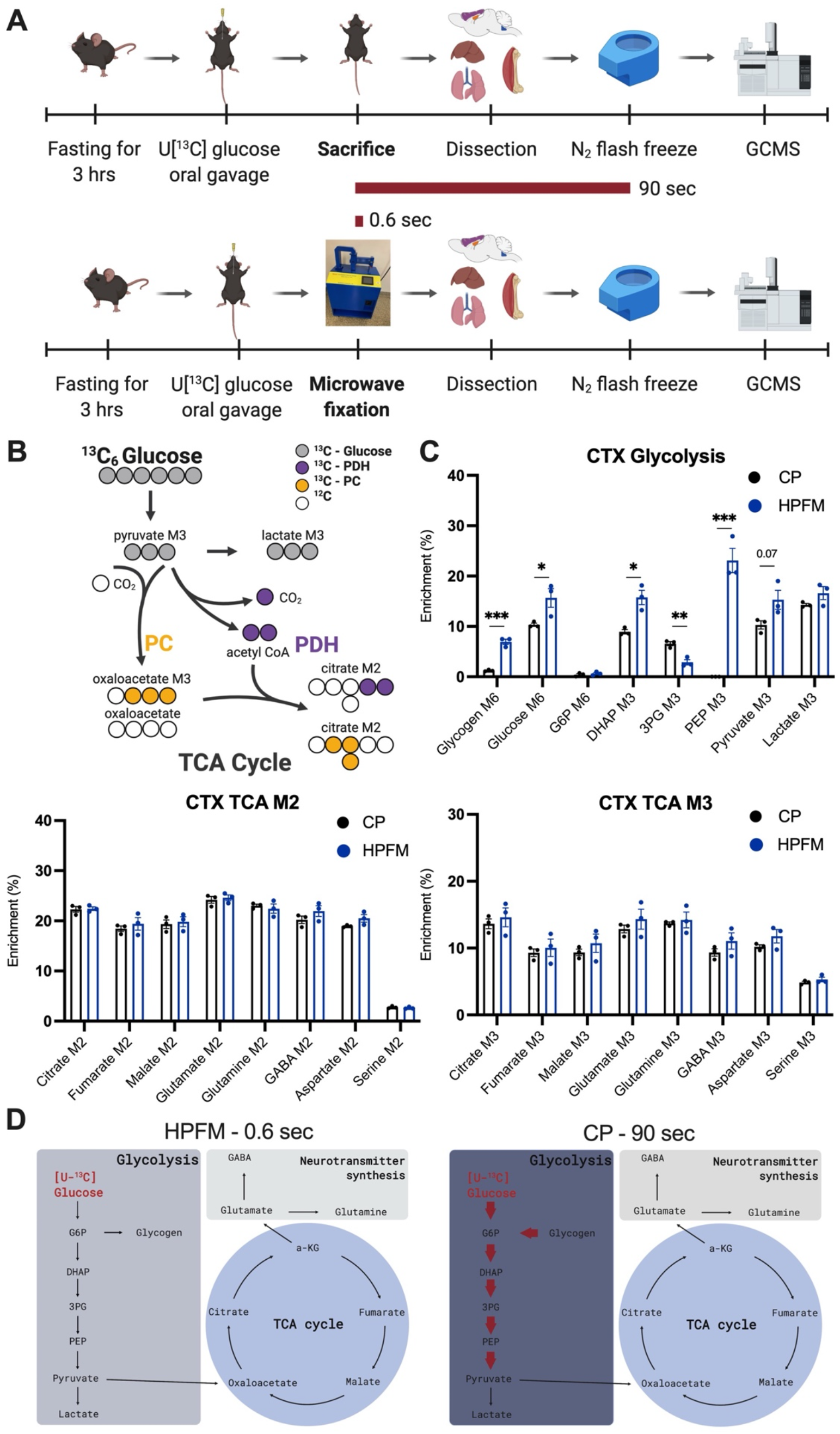
*In situ* high-power focused microwave fixation (HPFM) is compatible with stable isotope labeling. (**A**) Schematic of the experimental design. In a parallel experiment as shown in **Figure 1**, 6-week-old mice were fasted for 3 hours, gavaged with ^13^C_6_-glucose, euthanized by decapitation (*top*) or microwave fixation (*bottom*), brain regions were then dissected, flash-frozen, and analyzed by GCMS. (**B**) Schematic showing the stable isotope tracing glucose in glycolysis and TCA cycle and subsequent interpretations. (**C**) Representative isotopologue for M6 of glycogen, glucose, and G6P, M2 of citrate, fumarate, malate, glutamate, glutamine, GABA, aspartate, and serine, M3 of DHAP, 3PG, PEP, pyruvate, lactate, citrate, fumarate, malate, glutamate, glutamine, GABA, aspartate and serine within the neocortex regions between mice subjected to CP and HPFM. Values are presented as mean ±SEM (n=3 biological replicates), *p<0.05, **p<0.01, ***p<0.001, #p<0.0001, analyzed by student t-test. (**D**) Model of changes in glucose utilization within neocortex between mice euthanized by decapitation and microwave fixation. Bolded red arrows represent overutilized pathways during CP. G6P: glucose 6-phosphate, DHAP: dihydroxyacetone phosphate, 3PG:3-phosphoglyceric acid, PEP: phosphoenolpyruvate.

### Pooled metabolomics analysis of brain in a mouse model of T1DM using focused microwave

T1DM is a peripheral disease that is characterized by the lack of insulin production and persistent high circulating blood glucose. It is well-documented that T1DM negatively impacts neurological outcomes, even showing a relationship with Alzheimer’s related dementia^49,50^. To define the impact of T1DM on brain glucose metabolism using focused microwave, we utilized the STZ model of insulin impairment and hyperglycemia in mice and performed untargeted analysis of metabolite pools using GCMS. Mice were randomly assigned to two groups (n=8), one that was injected intraperitoneally (IP) with vehicle (citric acid), and another with a single dose of STZ to induce hyperglycemia for 14 days to mimic T1DM (Fig 3A-B). Both cohorts of mice were euthanized and fixed by focused microwave followed by surgical resection of multiple brain regions (DVC, HIPP, and CTX) as well as peripheral organs (muscle and liver). To our surprise, we observed increases only in glucose, glycogen, glucose 6-phosphate, and leucine in different brain regions (Fig. 3C-D, Table S5). The majority of the metabolite pools were relatively unchanged in the brain (Fig. 3C-D). These limited changes were not the case in peripheral organs, as we observed changes in multiple glycolytic and TCA cycle metabolites in the liver (Fig. S3, Table S6). Collectively our data suggest that STZ-induced hyperglycemia predominately impacts metabolite pools of peripheral organs but not the brain.

**Figure 3.**
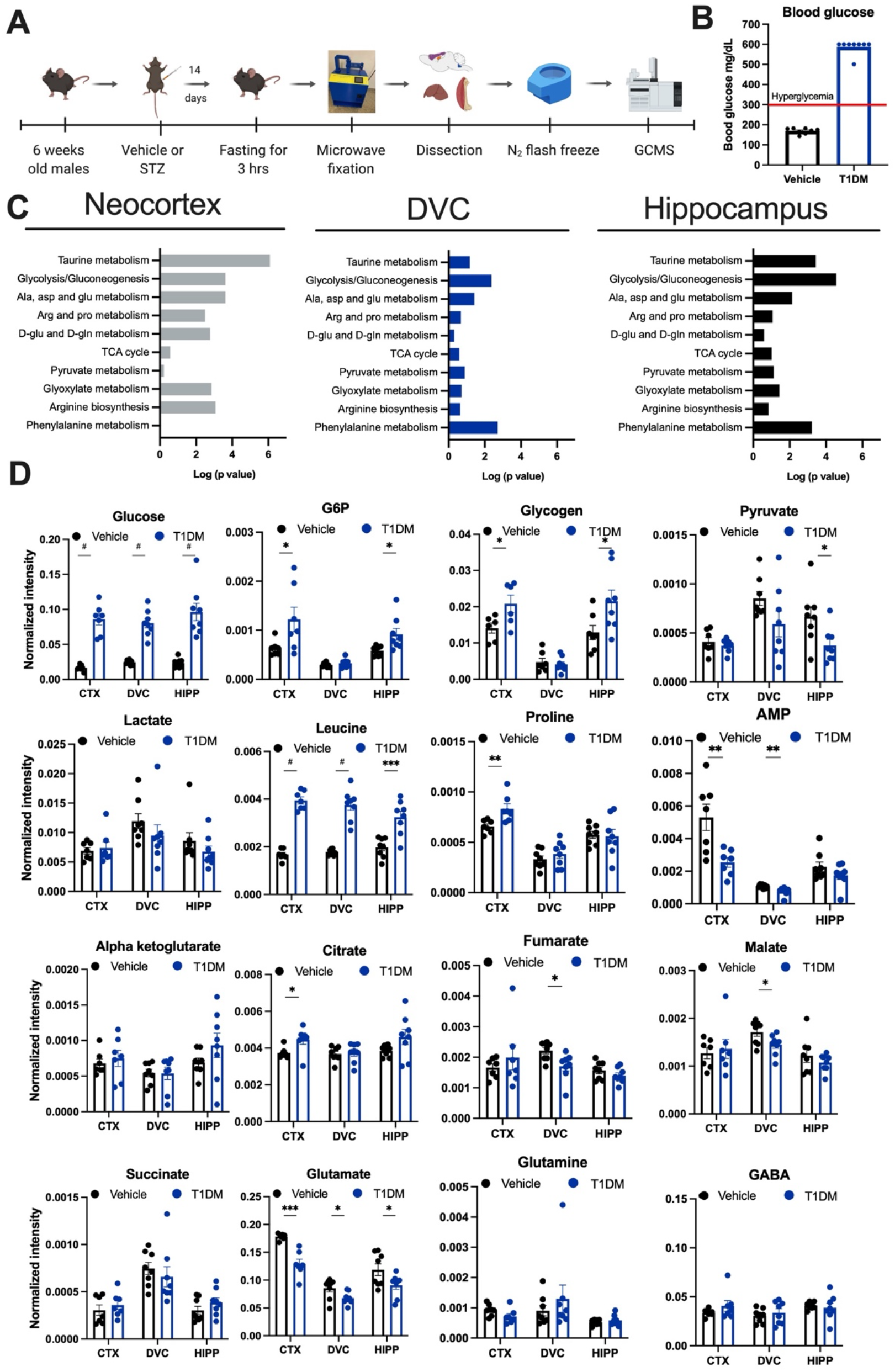
Pooled metabolomics analysis of mouse brain from a model of T1DM using focused microwave. (**A**) Schematic of the experimental design. 6-week-old mice were injected with either vehicle or 200mg/kg of STZ to induce hyperglycemia. After 14 days mice were fasted for 3 hours and euthanized by focused microwave. Brain regions were then dissected, flash-frozen, and analyzed by GCMS. (**B**) Blood glucose levels measured by hand-held glucometer in vehicle and T1DM mice after 14 days of hyperglycemia (n=8 biological replicates). (**C**) Pathway analysis (performed by Metaboanalyst) of CTX, DVC, and HIPP between vehicle treated and T1DM mice. (**D**) Representative metabolite levels from glycogen metabolism, glycolysis, and TCA cycle between vehicle and T1DM. Values are presented as mean ±SEM (n=6-8 biological replicates), *p<0.05, **p<0.01, ***p<0.001, #p<0.0001, analyzed by student t-test.

### Tracing ^13^C-glucose metabolism in the brain of a T1DM mouse model using focused microwave

Metabolic pathways are channeled within a cell and metabolite pools can be replenished through different metabolic substrates^7^. The TCA cycle metabolite pools can be replenished from glucose, glutamine, and fatty acid metabolism. Therefore, pooled metabolomics analysis does not fully illuminate unique substrate metabolism such as glucose. T1DM is primarily a disease of hyperglycemia, therefore it is crucial to define the glucose contribution to different metabolite pools especially when we did not observe major changes in the pooled analysis of STZ-treated mouse brains. To assess whether there are differences in glucose metabolism between vehicle and STZ treated mice we performed an additional animal experiment with vehicle and STZ-treated arms and performed stable isotope tracing of ^13^C_6_-glucose delivered through oral gavage with collection at either 30-minutes or 2-hours post-gavage (Fig 4A-B). Similar to above, all cohorts of mice were euthanized and fixed *in situ* with focused microwave followed by brain regional dissection and peripheral organ extraction. To our surprise, even though total pooled metabolites remain minimally affected in the STZ cohorts, we observed major decreases in glucose enrichment in central carbon metabolites and *de novo* synthesized amino acids. This phenotype is consistent across the CTX, HIPP, and the DVC for both time points (Fig. 4C-E, Fig. S4, Table S7) as well as in peripheral organs (Fig. S5, Table S8). Further, M2 and M3 isotopologues of citrate, malate, fumarate, glutamine, glutamate, and aspartate all exhibited a decrease in the STZ arm compared to the vehicle at both time points (Fig. 4C-E, Fig. S4). This result suggests down-regulation of pyruvate dehydrogenase and pyruvate carboxylase activity in STZ-induced hyperglycemia (i.e., T1DM) in multiple brain regions. Strikingly/Interestingly, when comparing the CTX to the DVC, we observed decreased lactate M3 isotopologue enrichment and increased M2 isotopologue of citrate, malate, and fumarate in the STZ-arm (Fig. 4F). These data could be interpreted as the degree of glucose hypometabolism being different among different brain regions (Fig. 4G).

**Figure 4.**
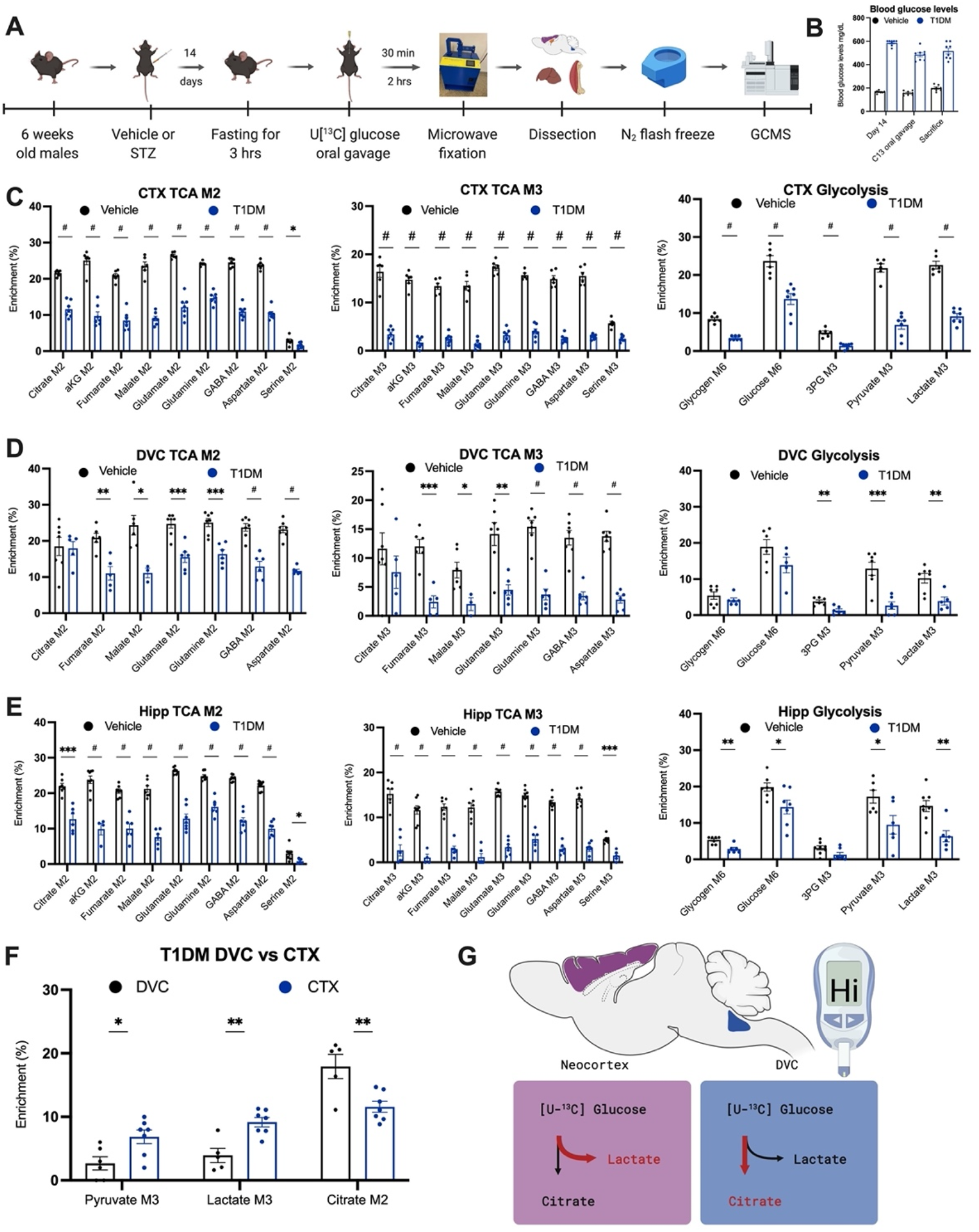
Stable isotope labeling in the brain of a mouse model of T1DM using focused microwave. (**A**) Schematic of the experimental design. 6-week-old mice were injected with either vehicle or 200mg/kg of streptozotocin (STZ) to induce hyperglycemia. After 14 days mice were fasted for 3 hours, gavaged with ^13^C_6_-glucose and euthanized by focused microwave. Brain regions were then dissected, flash-frozen, and analyzed by GCMS. (**B**) Blood glucose levels were measured by handheld glucometer before fasting, before tracer gavage delivery, and before microwave fixation (n=8). (**C-E**) Representative isotopologue for M6 of glycogen, glucose, and, M2 of citrate, aKG, fumarate, malate, glutamate, glutamine, GABA, aspartate and serine, and M3 of 3PG, pyruvate, lactate, citrate, aKG, fumarate, malate, glutamate, glutamine, GABA, aspartate and serine for three brain regions; CTX (**C**), DVC (**D**), and HIPP (**E**) of T1DM mice. Values are presented as mean ±SEM (n=6-8 biological replicates), *p<0.05, **p<0.01, ***p<0.001, #p<0.0001, analyzed by student t-test (**F**) M3 isotopologues of pyruvate and lactate and M2 isotopologue of citrate between CTX and DVC in T1DM. Values are presented as mean ±SEM (n=6-7 biological replicates), *p<0.05, **p<0.01, ***p<0.001, #p<0.0001, analyzed by student t-test. (**G**) Model of changes in glucose utilization in CTX and DVC and in T1DM. 3PG:3-phosphoglyceric acid, aKG: alpha-ketoglutarate.

### Downregulation of GLUT2, PDH, and PC proteins in unique neuronal cell layers

The power of stable isotopic tracing is the direct assessment of metabolic enzyme activities, through fractional enrichment, i.e., M2 and M3 isotopologues of citrate represent PDH and PC activities, respectively. Mice with STZ-induced hyperglycemia exhibit decreased glucose enrichment in central carbon metabolites and reduced M2 and M3 isotopologues of citrate (Fig 4C), therefore we continued to evaluate glucose transporters, PDH, and PC protein expression in cortical and HIPP regions using immunofluorescence (IF). First, we performed IF for GLUT1, GLUT2, and GLUT3 glucose transporters, which have been shown to be affected by hyperglycemia in peripheral tissues^51–54^. We did not observe significant changes in percent positive area in GLUT1 and GLUT3 expression in the different brain regions (Fig S6), however GLUT2, a neuronal glucose transporter displays a 15% to 70% decrease of percent positive areas across the CTX, HIPP, and DVC (Fig. 5A). We observed a similar decrease in PDH and PC protein by IF in cortical regions and the DVC (Figure 5B and Fig. S6). Interestingly, CA3 neuronal layer of HIPP was the only sub-region affected by hyperglycemia, while CA1 and CA2 layers exhibited no differences in protein expression between PC and PDH assessed by IF (Figure 5B and Fig. S6). Collectively, these data suggest STZ-induced hyperglycemia results in downregulation of metabolic enzymes at the protein level that correlate with glucose hypometabolism of the brain.

**Figure 5.**
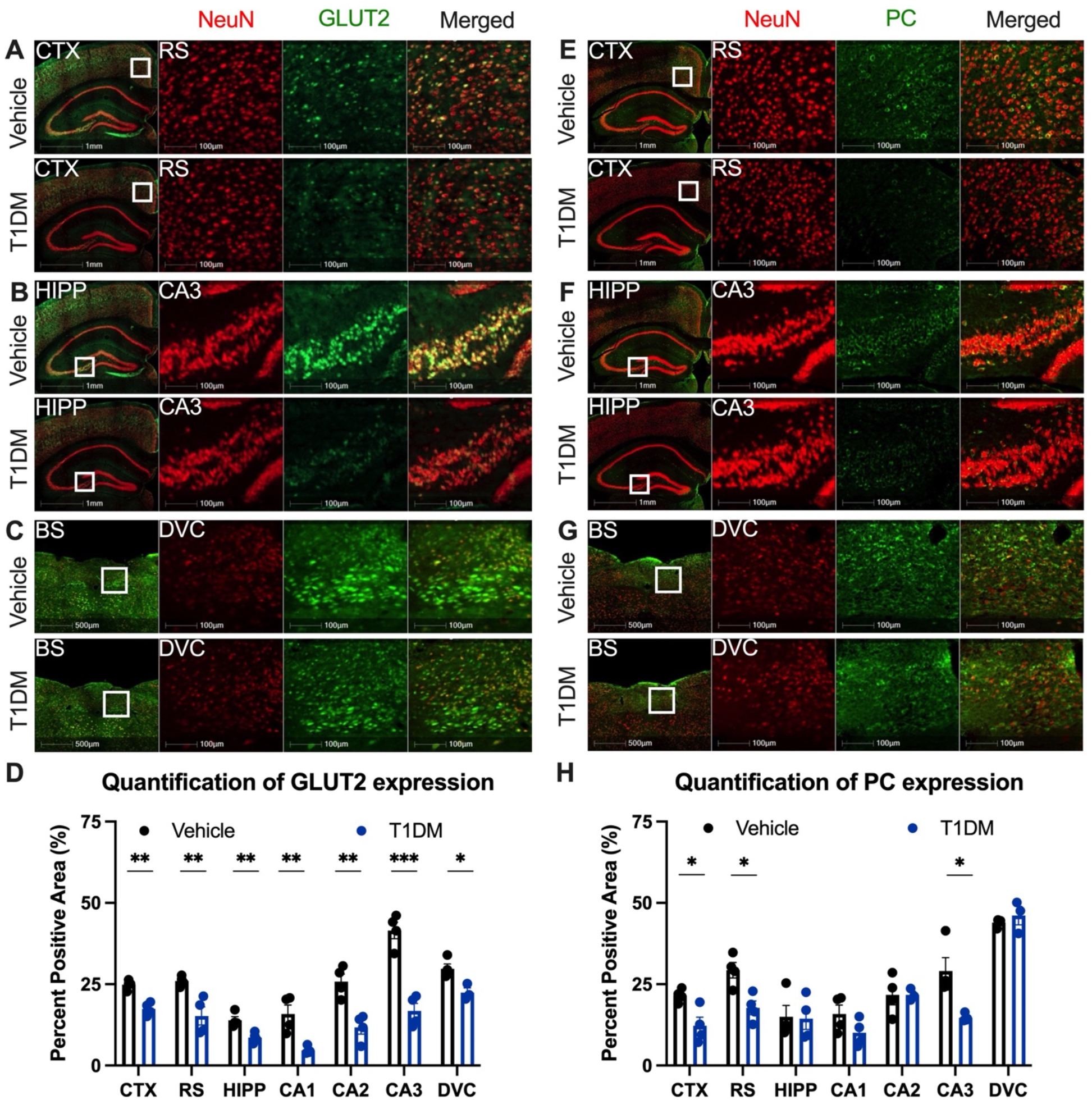
Immunofluorescent analysis of GLUT2 and PC expression in CTX, HIPP and brainstem (BS) in a mouse model of T1DM. (**A-D**) 20μm coronal sections of the mouse brain were stained with GLUT2 (green) and NeuN (red) and quantified in in the CTX (**A**), HIPP (**B**), and BS (**C**). Zoomed-in images are shown for the retro-splenial cortex (RS), Cornu Ammonis 3 (CA3) of HIPP, and DVC of BS. (**D**) Quantification of % positive pixels for GLUT2 using HALO software, area of interest include whole neocortex, RS (zoomed in), whole HIPP, CA1-CA3 (zoomed in), and DVC following mouse brain atlas, and quantification was run for positive pixels in each channel and colocalization. Values are presented as mean ±SEM (n=4 biological replicates), *p<0.05, **p<0.01, ***p<0.001, analyzed by student t-test. (**E-H**) 20 μm coronal sections of the mouse brain were stained with pyruvate carboxylase (PC) (green) and NeuN (red) and quantified in in the CTX (**E**), HIPP (**F**), and BS (**G**). Zoomed in image are shown for the RS, CA3 of HIPP, and DVC of BS. (**H**) Quantification of % positive pixels for PC using HALO software, area of interest including whole CTX, RS (zoomed in), whole HIPP, CA1-CA3 (zoomed in), and DVC following mouse brain atlas, and quantification was run for positive pixels in each channel and colocalization. Values are presented as mean ±SEM (n=4 biological replicates), *p<0.05, **p<0.01, ***p<0.001, analyzed by student t-test. Scale bars are either 1mm, 500μm, or 100μm. PC: pyruvate carboxylase, GLUT2: glucose transporter 2.

## DISCUSSION

There is a major scientific interest in deepening our understanding of the brain metabolome ^55,56^, as many speculate it will aid in our understanding of brain physiology and could be key to understanding neurological disease pathology. Brain metabolism is a highly heterogeneous and rapid cycling series of events, therefore capturing an accurate snapshot of the brain metabolome is technologically challenging. In this study, we employed focused microwave to create a snapshot of the brain metabolome through *in situ* heat inactivation and fixation of metabolic enzymes. By comparing focused microwave fixation and conventional cryo-preservation, we found that focused microwave significantly retained total pools of glycogen, glucose, and citrate, but showed decreased levels of lactate, malate, and succinate. Further, we applied focused microwave to study changes in brain metabolism in a STZ-induced mouse model of T1DM and observed profound glucose hypometabolism in multiple brain regions of diabetic mice subject to hyperglycemia. Our study highlighted the importance of capturing the accurate brain metabolome and supports the use of focused microwave to study brain metabolism. When focused microwave is not available, data interpretation should be carefully examined with the assumption that non-physiological metabolism occurs during cryo-preservation.

Currently, strategies to study brain metabolism include total pooled metabolomics analyses for large-scale pathway coverage^26^ and in certain cases, stable isotope tracing is employed to study metabolic flux or enzyme activity^21,57^. We observed a number of interesting differences between focused microwave and cryo-preservation in total pooled metabolomics analysis. Primarily, we observed drastic increases in glycogen, glucose, citrate, and aKG, but a decrease in lactate, succinate, and malate in mouse brain fixed *in situ* with focused microwave, versus cryo-preservation. These data suggest increased utilization of glycogen and glucose for glycolysis and lactate production, as well as TCA cycle anaplerosis during the cryo-preservation process. Interestingly, the metabolic shift toward lactate production and anaplerosis are often observed during hypoxia^58^ and occur frequently during cancer metabolism^59^. One possible explanation of this metabolic phenotype is that after euthanasia, the lack of circulation supplying the brain creates a hypoxic environment for the metabolically active brain cells, resulting in a shift towards a hypoxic metabolic phenotype. It is worth noting we did not observe a difference in amino acid abundances between focused microwave and cryo-preservation. Amino acids such as glutamate, glutamine, aspartate, or GABA remain similar between the two different brain harvest strategies.

Stable isotope tracing is a powerful technique to study metabolic flux and enzyme activity in vivo and is frequently used to study brain metabolism^21,57^. We performed an isotopic tracing experiment using an *in vivo* oral delivery method developed previously^44^ to test the effects of *in situ* fixation by focused microwave on stable isotope enrichment of central carbon metabolites. In line with pooled metabolomics analysis, we observed evidence of increased glucose and glycogen utilization demonstrated by decreased M6 isotopologue enrichment of glucose and glycogen, and increased M3 isotopologue enrichment of 3-PG during cryopreservation. To our surprise, we did not observe major changes in isotopic enrichment of TCA cycle metabolites. M2 and M3 isotopologues of citrate, fumarate, and malate were consistent between focused microwave and cryo-preservation. These data imply that although the total pool of metabolites was changing during cryo-preservation, it did not affect the isotopic ratios or enrichment. Further, both labeled and unlabeled ^13^C fractions were either produced or consumed at the same rate. Based on these data, we recommend the use of stable isotope tracing to study brain metabolism when focused microwave is not available.

T1DM affects about 20 million people worldwide and chronic hyperglycemia leads to micro/macro-vascular diseases, retinopathy, peripheral neuropathy, and liver disease^60^. Recent reports have highlighted the impact of T1DM on brain glucose metabolism and T1DM patients have increased risk of developing late onset dementia^61^. We interrogated the changes in pooled brain metabolome and more specifically, glucose metabolism using ^13^C_6_-glucose isotopic tracing in the STZ treated mouse model of T1DM by focused microwave. We did not observe major changes in pooled metabolites except for glucose and glycogen which remained higher in the vehicle cohort. We then performed ^13^C-glucose tracing through oral gavage. We observed glucose hypometabolism in all three regions we examined. Decreased enrichment of glycolytic and TCA cycle isotopologues were noted in multiple regions of the brain including CTX, HIPP, and DVC. These data are in agreement with previous reports showing brain glucose hypometabolism in T1DM patients^62^, impaired hepatic glycogen accumulation^63–66^, glycogenolysis^64,67^ and increased hepatic glucose production^68–71^. Further, while glucose metabolism is decreased, total pools of metabolites remained unchanged, suggesting alternative fuel sources are being utilized by the brain in T1DM mice that warrant further investigation. Notably, glucose metabolism differed between the CTX and the DVC within the T1DM cohort. Within the DVC, differences in isotopic enrichment of citrate and lactate suggest glucose is preferentially oxidized via the TCA cycle in hyperglycemic mice. It is worth noting that the blood brain barrier is permeable throughout much of the DVC and the region could have alternative metabolic controls compared to the CTX and HIPP.

### Limitations and future directions

In this study, we aimed to improve the assessment of the brain metabolome by decreasing the post-mortem impact following brain resection. We would like to highlight that rodent brain metabolism is highly sensitive to stimuli such as handling and stress^72^, anesthesia^73^, and time post-euthanasia^29^. Additional steps will be needed to control these factors in the future to truly capture the “normal” brain metabolome. Another limitation of this study is the utilization of STZ induced model of T1DM, as direct injection of STZ to the brain has been shown to induce neuroinflammation and alter brain physiology^74^. Additional diabetes mouse models or human patients should be evaluated in the future to confirm the glucose hypometabolism phenotype. Future studies should focus on isolating single cell types through fluoresce-activated cell sorting (FACS) or magnetic separation methodologies post focused microwave, to define the cell type specific brain metabolism in healthy and diseased tissue. It would also be of interest to test whether focused microwave is compatible with new single-cell technologies such as matrix-assisted laser desorption ionization MALDI mass spectrometry imaging^17^, single cell RNAseq^75^, and spatial proteomics analyses^76,77^.

## Materials and Methods

### Chemicals and Reagents

HPLC grade methanol (34860-4X4L-R) and methoxyamine hydrochloride (226904-25G) were purchased from Sigma Aldrich (Burlington, MA, USA). Silylation Reagents, MSTFA + 1% TMCS Reagent (TS-48915) and pyridine (TS-27530) were purchased from Thermo Fisher Scientific (Waltham, MA, USA). Amber vials (5184-3554), blue screw caps (5182-0717) were purchased from Agilent Technologies (Santa Clara, CA, USA). Streptozotocin (572201), PBS (P3813), Triton X-100 (X100), Tween 20 (P1379) were purchased from Sigma-Aldrich (St. Louis MO). Sodium citrate dihydrate (BP327), Citric acid anhydrous (A940) normal goat serum (31873) and OCT compound (23-730-571), 2-methylbutane (019387AP), and ProlongGlass (P36984) from Thermo Fisher Scientific. [U-^13^C] Glucose was obtained from Cambridge Isotope Laboratories (Tewksbury, MA, USA).

### Animals

All experiments were performed on either male Vgat-ires-Cre knock-in mice (Slc32a1tm2(cre)lowl/J; 016962; The Jackson Laboratory, Bar Harbor, ME) or C57BL6 background (000664; The Jackson Laboratory, Bar Harbor, ME). Mice were housed and cared for in the University of Kentucky Division of Laboratory Animal Resources facilities under normal 14:10 light-dark condition with food (Tekad 2018, Indianapolis, IN) and water available ad libitum, except where noted and with accordance to protocols approved by the University of Kentucky Animal Care and Use Committee.

### Induction of hyperglycemia and in vivo glucose assessments

At 6 weeks of age, mice were injected with streptozotocin (STZ; 200mg/kg; i.p.; Sigma-Aldrich, St. Louis, MO) in citric acid (CA, 0.1M), after a 6 hour fast. Mice were monitored daily for weight and blood glucose levels. Blood glucose levels were measured by handheld glucometer (Nova Max Plus; received from American Diabetes Wholesale (ADW), Pompano Beach, FL), using 0.3μL of blood from the tail vein. Nova Max glucose test strips were used (#8548043523, ADW). Blood concentration above 300mg/dL was considered hyperglycemic. Mice were used for experiments after at least 14 days (14-16 days) of hyperglycemia.

### Gavage of [U-^13^C] glucose

[U-^13^C] Glucose (Cambridge Isotope Laboratories, Tewksbury, MA, USA) was dissolved in ddH2O (Millipore Milli-Q, Bedford, MA, USA) based on the average mouse cohort bodyweight (2 g [U-^13^C] glucose/kg bodyweight). After 3 hours fast, 250 μL of glucose solution was administered via oral gavage. Tissues were collected at 30 minutes and 2 hours. To assess the blood glucose levels before fasting, at tracer delivery and before microwave fixation handheld glucometer (Nova Max Plus; received from American Diabetes Wholesale, Pompano Beach, FL) was used.

### Microwave fixation

Mice were euthanized by microwave fixation system at 5kH for 0.6 seconds (MMW-05, Muromachi Kikai Company, Japan). Brain regions (HIPP, CTX, DVC), as well as muscle and liver, were dissected postmortem.

### Immunofluorescence

Mice were transcardially perfused with Heparin-PBS (10 units/mL), followed by 4% PFA. Tissue was post-fixed for 24 hours in 4%PFA and then cryoprotected with 30% sucrose in 0.01M PBS. The tissue was frozen with isopentanes and cut at 20μm with a sliding microtome. Staining was done free-floating in 24-well plates without inserts. Tissue was rinsed with 0.01M PBS, incubated for 1 hour in 5% normal goat serum in 0.3% Triton X-100 in PBS. The tissue was then incubated at room temperature in the primary antibody (table 1) at diluted in 1% normal goat serum, followed by 3 x 15 minutes rinses in 0.05% Tween-20 in PBS. Tissue was then incubated for 1 hour in secondary antibody diluted in 1% normal goat serum in PBS. For multiple labeling, primary antibodies were incubated for 2 hours, and secondary antibodies for 1 hour. Tissue was then exposed to DAPI for 5 minutes, mounted on the slides and cover slipped with Prolong Glass. Digital images were acquired through the Zeiss Axio Scan Z.7 digital slide scanner at 20X magnification and 12 Z-stacks. Figures were captured using HALO software (v3.3.2541.345, Indica Labs, Albuquerque, NM). Analysis was done by outlining the regions of interest in the HALO software and quantifying the positive pixels.

| **Primary antibodies** | **Host** | **Company and catalog #** | **Dilution** | **Conditions** |
| --- | --- | --- | --- | --- |
| GLUT1 | Rabbit | Abcam ab115730 | 1:200 | Overnight, RT |
| GLUT2 | Rabbit | Novus NBP2-22218 | 1:200 | Overnight, RT |
| GLUT3 | Rabbit | Alomone AGT-023 | 1:200 | Overnight, RT |
| Pyruvate Carboxylase | Rabbit | GeneTex GTX121987 | 1:200 | Overnight, RT |
| Pyruvate Dehydrogenase (ser 293) | Rabbit | EMD Millipore AP1062 | 1:200 | Overnight, RT |
| NeuN | Mouse | Abcam ab279295 | 1:1000 | 1 hour, RT |
| NeuN | Rabbit | Abcam ab236870 | 1:1000 | 1 hour, RT |
| GFAP | Chicken | GeneTex GTX85454 | 1:2000 | 2 hour, RT |
| DAPI |  | Novus NBP2-31156 | 1:5000 | 5 minutes, RT |

| Secondary antibodies | Fluorophore | Company and catalog # | Dilution | Conditions |
| --- | --- | --- | --- | --- |
| Goat anti-chicken | Alexa 594 | A-11042 | 1:400 | 1 hour, RT |
| Goat anti-rabbit | Alexa 488 | A-11034 | 1:400 | 1 hour, RT |
| Goat anti-mouse | Alexa 647 | A-21236 | 1:400 | 1 hour, RT |

### Sample Preparation

Brains were removed immediately post-mortem, and washed once with PBS, twice with diH_2_O, blotted dry, and snap frozen in liquid nitrogen. Other set of brains were snap frozen after microwave fixation as described above. The frozen tissues were pulverized to 10 μm particles in liquid N_2_ using a Freezer/Mill Cryogenic Grinder (SPEX SamplePrep). Brain regions were extracted with 1ml of 50% methanol in the grinder, while for muscle and liver twenty milligrams of each pulverized tissue were extracted in 1ml of 50% methanol and separated into polar (aqueous layer), and protein/DNA/RNA/glycogen pellet. The polar fraction was dried at 10^-3^ mBar using a SpeedVac (Thermo) followed by derivatization.

### Pellet Hydrolysis

Hydrolysis of the protein/DNA/RNA/glycogen pellet was performed by first resuspending the dried pellet in diH2O followed by the addition of equal parts 2N HCl. Samples were vortexed thoroughly and incubated at 95°C for 2 hours. The reaction was quenched with 100% methanol with 40μM L-norvaline (as an internal control). The sample was incubated on ice for 30 minutes and the supernatant was collected by centrifugation at 15,000rpm at 4°C for 10 minutes. The collected supernatant was subsequently dried for 30 minutes by vacuum centrifuge at 10^-3^ mBar.

### Sample Derivatization

Dried polar and glycogen samples were derivatized by the addition of 20mg/ml methoxyamine hydrochloride in pyridine and sequential addition of N-methyl-trimethylsilyl-trifluoroacetamide (MSTFA). Both reactions were incubated for 60 minutes at 60°C with thorough mixing in between addition of solvents. The mixture was then transferred to a v-shaped amber glass chromatography vial and analyzed by GCMS.

### GCMS Quantification

GCMS protocols were similar to those described previously^78,79^, except a modified temperature gradient was used for GC: Initial temperature was 130°C, held for 4 minutes, rising at 6°C/minutes to 243°C, rising at 60°C/minutes to 280°C, held for 2 minutes. The electron ionization (EI) energy was set to 70 eV. Scan (m/z:50-800) and full scan mode were used for metabolomics analysis. Mass spectra were translated to relative metabolite abundance using the Automated Mass Spectral Deconvolution and Identification System (AMDIS) software matched to the FiehnLib metabolomics library (available through Agilent) for retention time and fragmentation pattern matching with a confidence score of > 80^80–82^. Data was further analyzed using the Data Extraction for Stable Isotope-labelled Metabolites (DEXSI) software package^83^. Untargeted metabolomics data was normalized to total ion chromatogram. For glucose tracer raw data was exported and correction for natural abundance was done by IsoCorrectoR^84^. Fractional enrichment of each metabolite was calculated as the relative abundance of each isotopologue relative to the sum of all other isotopologues. Mean enrichment was calculated as sum of fractional enrichment of labeled isotopologues (M1, M2, M3…). Untargeted metabolomics was performed as described in Young et al 2019. For principal component analysis, pathway impact analysis the online tool Metaboanlyst was used (https://www.metaboanalyst.ca/). Data was auto scaled and log transformed.

### Statistics

Statistcal analyses were carried out using GraphPad Prism. All numerical data are presented as mean ± SEM. Column analysis was performed using t-test. A P-value less than 0.05 was considered statistically significant. Statistical power and sample size calculations for animal studies were performed based on the current focused microwave data set (neocortex). A sample of n=3 / group would provide 80% power to detect a 3.1 standard deviation difference in the mean abundance level of a metabolomic feature between experimental groups based on a two-sample t-test at 5% significance level.

## Data and materials availability

All data are available in the main text or the supplementary materials. This study includes no data deposited in external repositories.

## Author contributions

Conceptualization, R.C.S., B.N.S., and J.A.J.; methodology, R.C.S., B.N.S., and J.A.J.; investigation, J.A.J., M.B.W., L.E.A.Y., K.H.M., T.R.H., M.D.B., K.E.B., P.T.C., M.I.W, L.PY.S., W.C.S., R.C.B., L.R.C., and C.W.; writing – original draft, R.C.S. and J.A.J.; writing – review & editing, R.C.S., J.A.J.; L.E.A.Y., L.R.C., T.R.H., B.N.S., and M.S.G.; funding acquisition, R.C.S., M.S.G., and B.N.S.; resources, R.C.S., M.S.G. and B.N.S.; supervision, R.C.S., M.S.G., and B.N.S.

## Acknowledgments

We would like to thank Dr. Binoy Joseph for assistance with the Z7 microscope, and Dr. Kathryn E Saatman for technical help with free-floating IF. We also thank Dr. Christopher Richards from UKs Light Microscopy Core for assistance. This study was supported by the National Institute of Health (NIH) grants R35 NS116824 (M.S.G.), P01 NS097197 (M.S.G.), NIH grant R01 AG066653 (R.C.S.), NIH grant R01 CA266004 (R.C.S.), NIH NIDDK R01 DK122811 (BNS), NIH NINDS R01 NS092552 (BNS), NIH/NCI F99CA264165 (L.E.A.Y.), NIH/NCI training grant T32CA165990 (L.R.C.), St Baldrick’s Career Development Award (R.C.S.), V-Scholar Grant (R.C.S.), and Rally Foundation Independent Investigator Grant to (R.C.S.).

---

**Figure S1.**
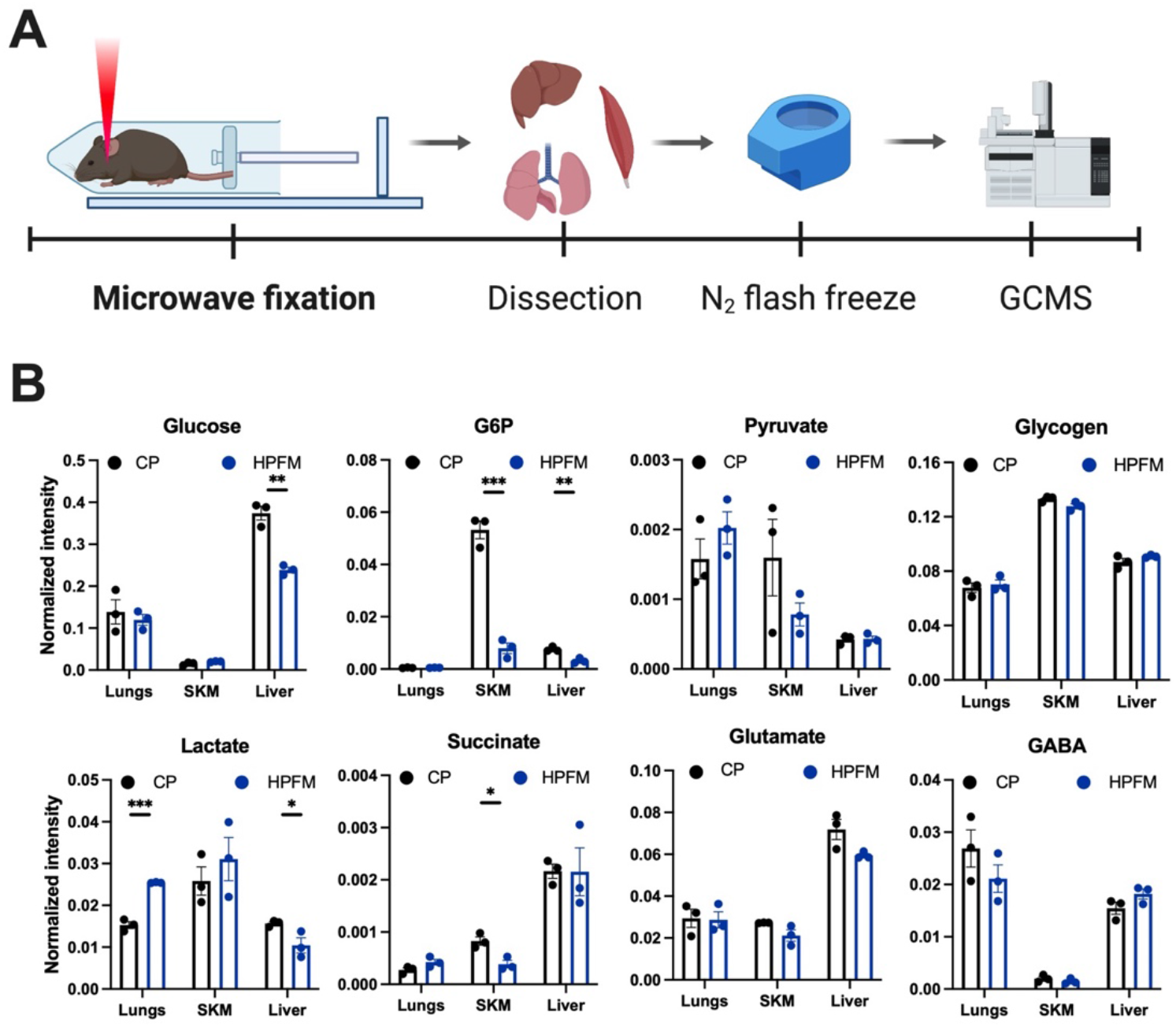
*In situ* high power focused microwave fixation (HPFM) to study metabolome in peripheral organs. (**A**) Schematic of the HPFM showing the energy beam hitting the brain but not peripheral organs. (**B**) Representative metabolite levels from glycogen metabolism, glycolysis, and TCA cycle between cryo-preservation (CP) and HPFM. Values are presented as mean ±SEM (n=3 biological replicates), *p<0.05, **p<0.01, ***p<0.001, analyzed by student t-test. G6P: glucose 6-phosphate.

**Figure S2.**
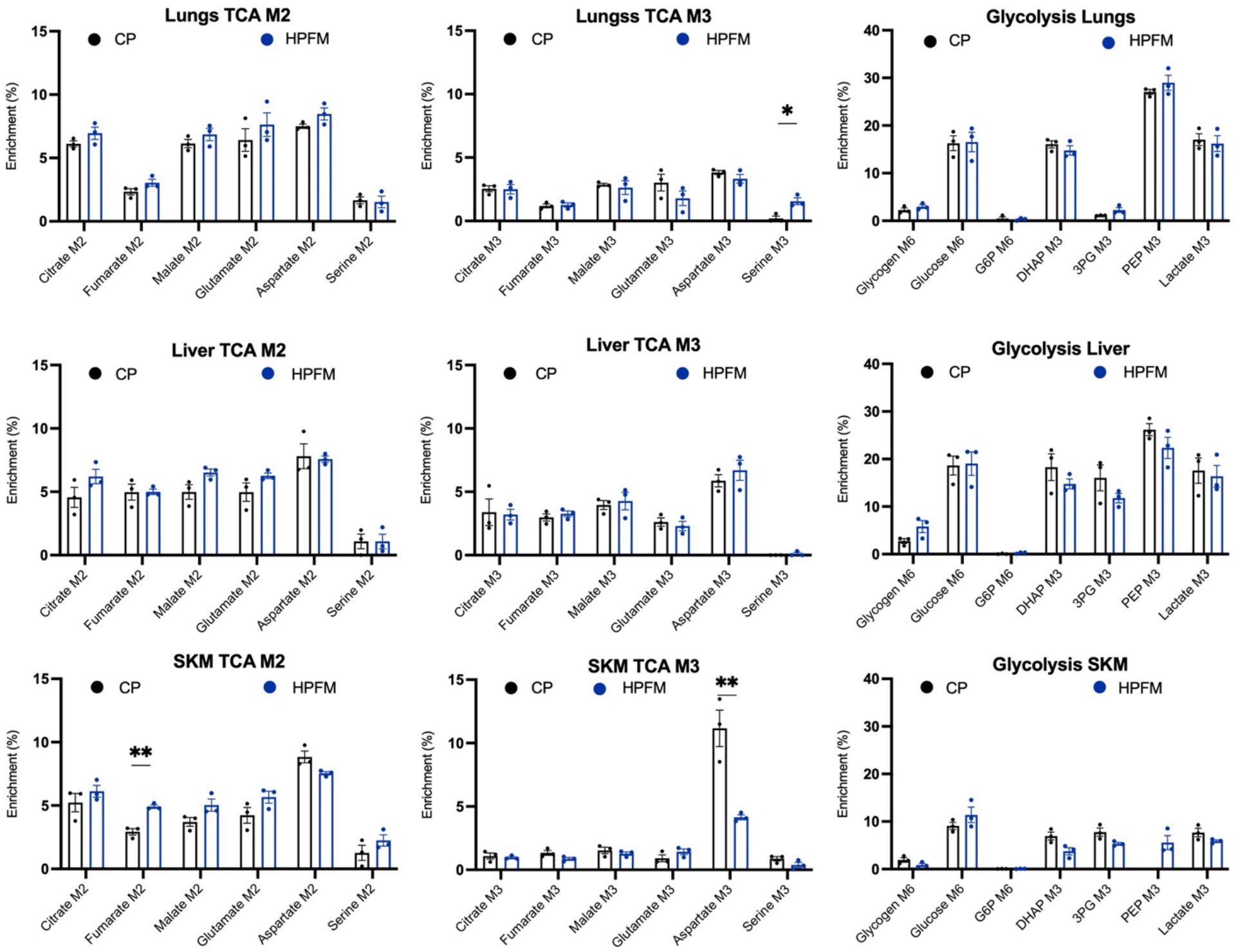
^13^C_6_ glucose isotope tracing does not alter in the lung, liver, or skeletal muscle (SKM) with high power focused microwave fixation (HPFM). Representative isotopologue for M6 of glycogen, glucose, and G6P, M2 of citrate, fumarate, malate, glutamate, aspartate, and serine, M3 of DHAP, 3PG, PEP, lactate, citrate, fumarate, malate, glutamate, aspartate and serine between mice subjected to CP and HPFM. Values are presented as mean ±SEM (n=3 biological replicates), *p<0.05, **p<0.01, analyzed by student t-test.

**Figure S3.**
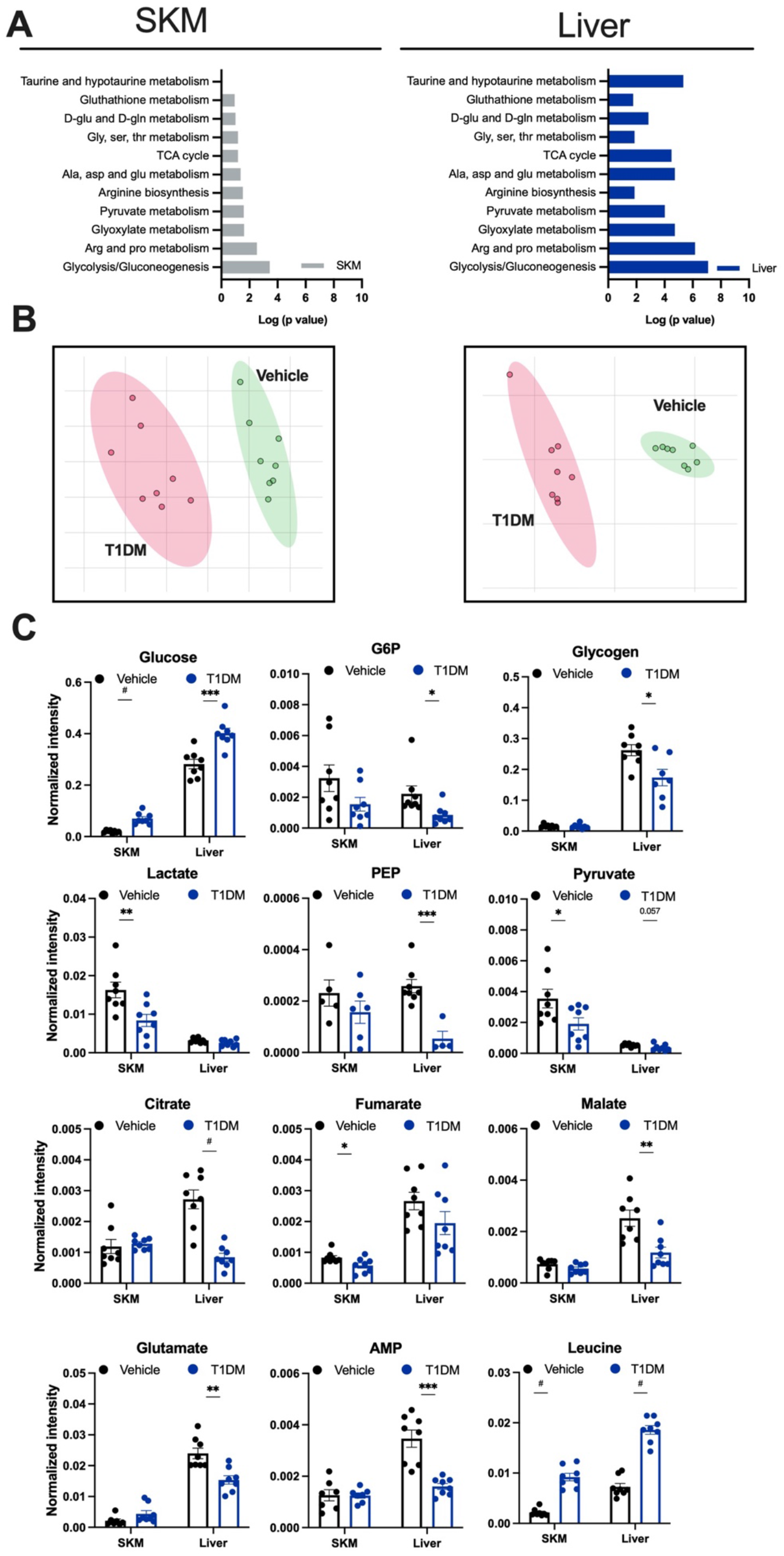
Pooled metabolomics analysis of skeletal muscle (SKM) and liver from a model of T1DM using focused microwave. (**A**) Pathway analysis (performed by Metaboanalyst) of SKM and liver between vehicle and T1DM. (**B**) Partial Least Squares-Discriminant Analysis (PLS-DA) of SKM and liver metabolites between vehicle and T1DM animals (n=6-8). (**C**) Representative metabolite levels from glycogen metabolism, glycolysis, and TCA cycle between vehicle and T1DM. Values are presented as mean ±SEM (n=6-8 biological replicates), *p<0.05, **p<0.01, ***p<0.001, #p<0.0001, analyzed by student t-test. G6P: glucose 6-phosphate, PEP: phosphoenolpyruvate, AMP: adenosine phosphate

**Figure S4.**
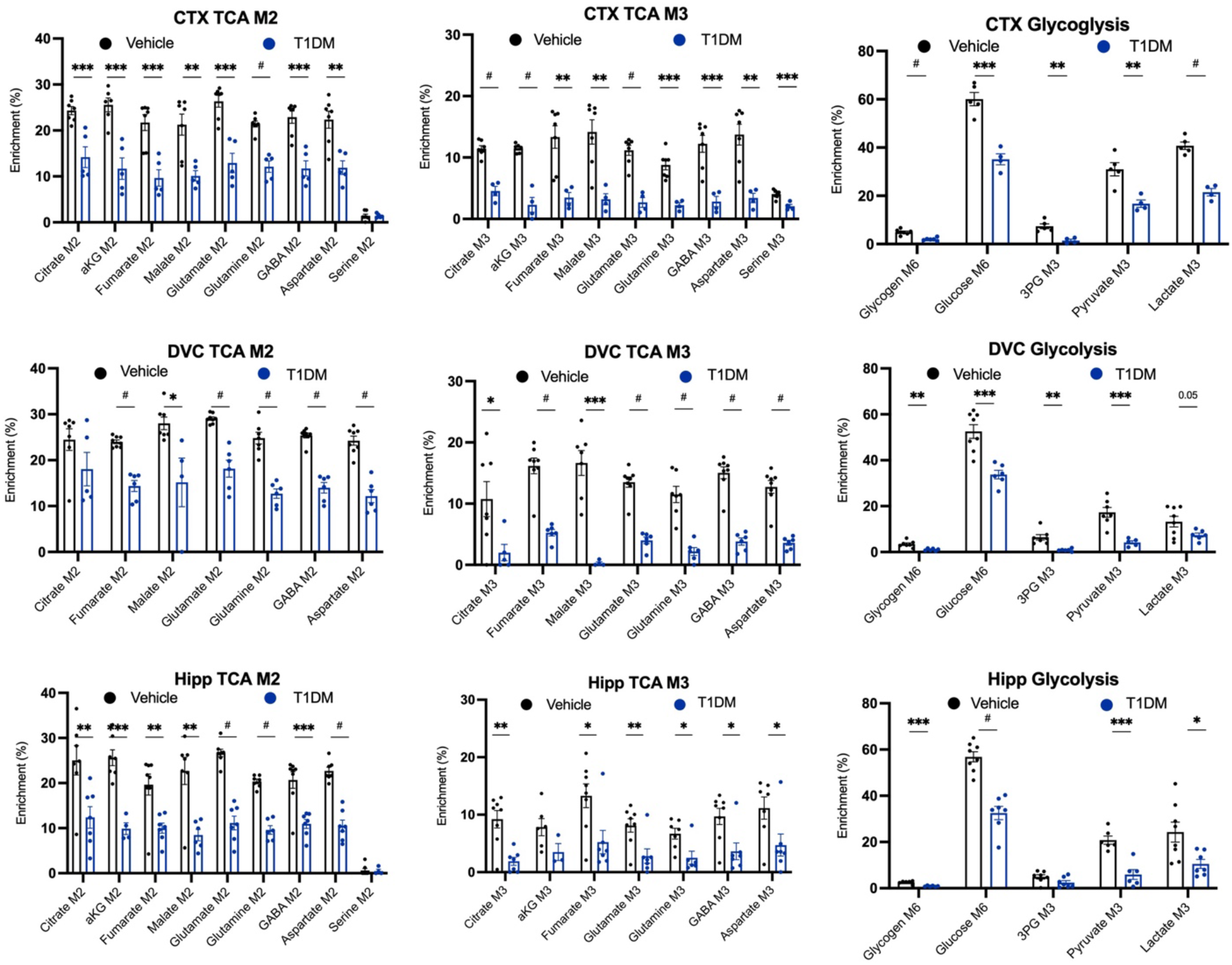
Stable isotope labeling in the brain of a mouse model of T1DM using HPFM at 30 min. Representative isotopologue for M6 of glycogen, glucose, and, M2 of citrate, aKG, fumarate, malate, glutamate, glutamine, GABA, aspartate and serine, and M3 of 3PG, pyruvate, lactate, citrate, aKG, fumarate, malate, glutamate, glutamine, GABA, aspartate and serine for three brain regions; neocortex (CTX), dorsal vagal complex (DVC) and hippocampus (HIPP) of T1DM animals. Values are presented as mean ±SEM (n=4-7 biological replicates), *p<0.05, **p<0.01, ***p<0.001, #p<0.0001, analyzed by student t-test. 3PG:3-phosphoglyceric acid, aKG: alpha-ketoglutarate.

**Figure S5.**
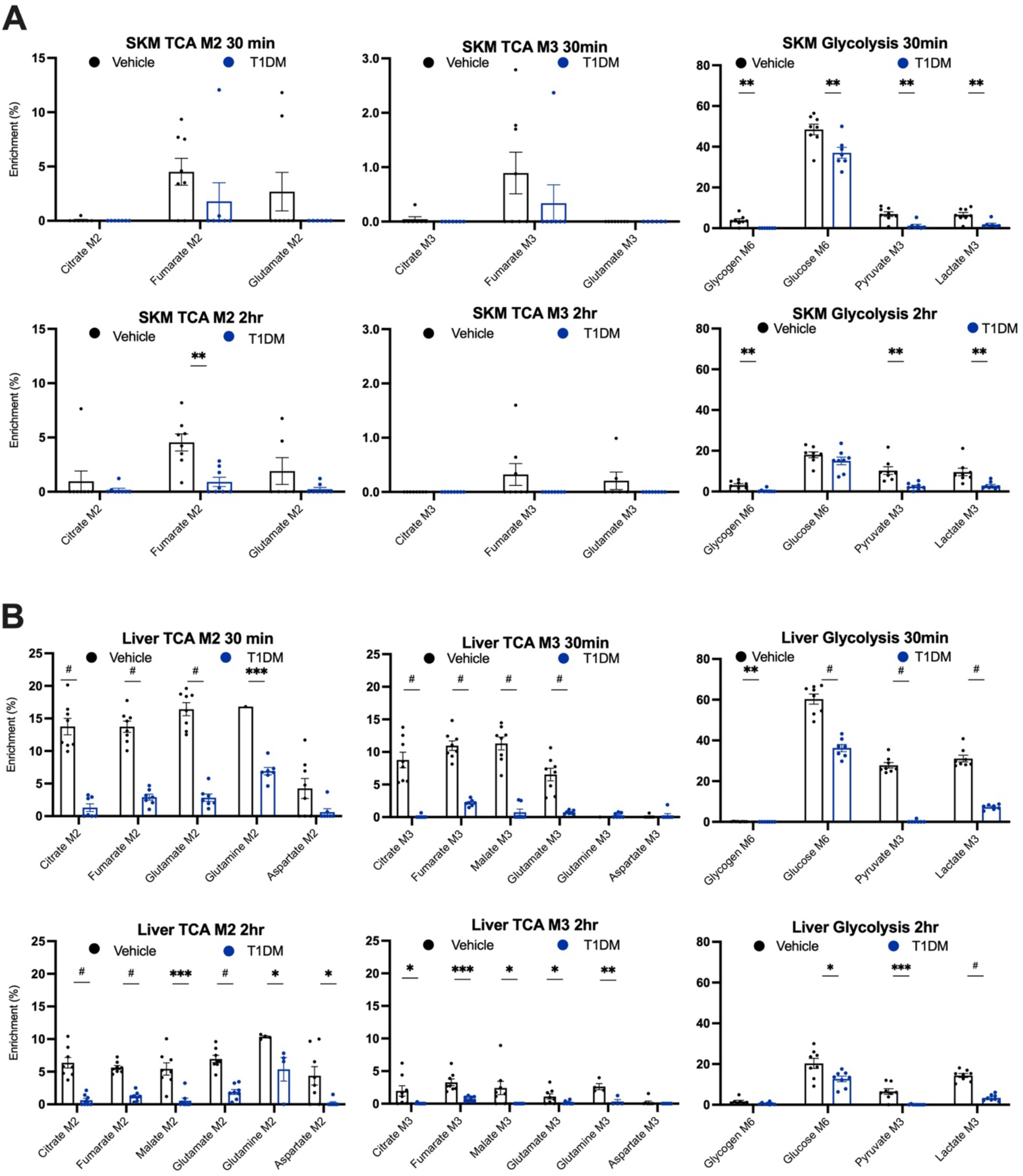
Stable isotope labeling at 30 min (**A**) and 2 hrs (**B**) in T1DM mice using focused microwave in skeletal muscle (SKM) and liver. Representative isotopologue for M6 of glycogen, glucose, and, M2 of citrate, fumarate, malate, glutamate, glutamine, and aspartate, and M3 of pyruvate, lactate, citrate, fumarate, malate, glutamate, glutamine, and aspartate for SKM and liver of T1DM animals. Values are presented as mean ±SEM (n=7-8 biological replicates), *p<0.05, **p<0.01, ***p<0.001, #p<0.0001, analyzed by student t-test.

**Figure S6.**
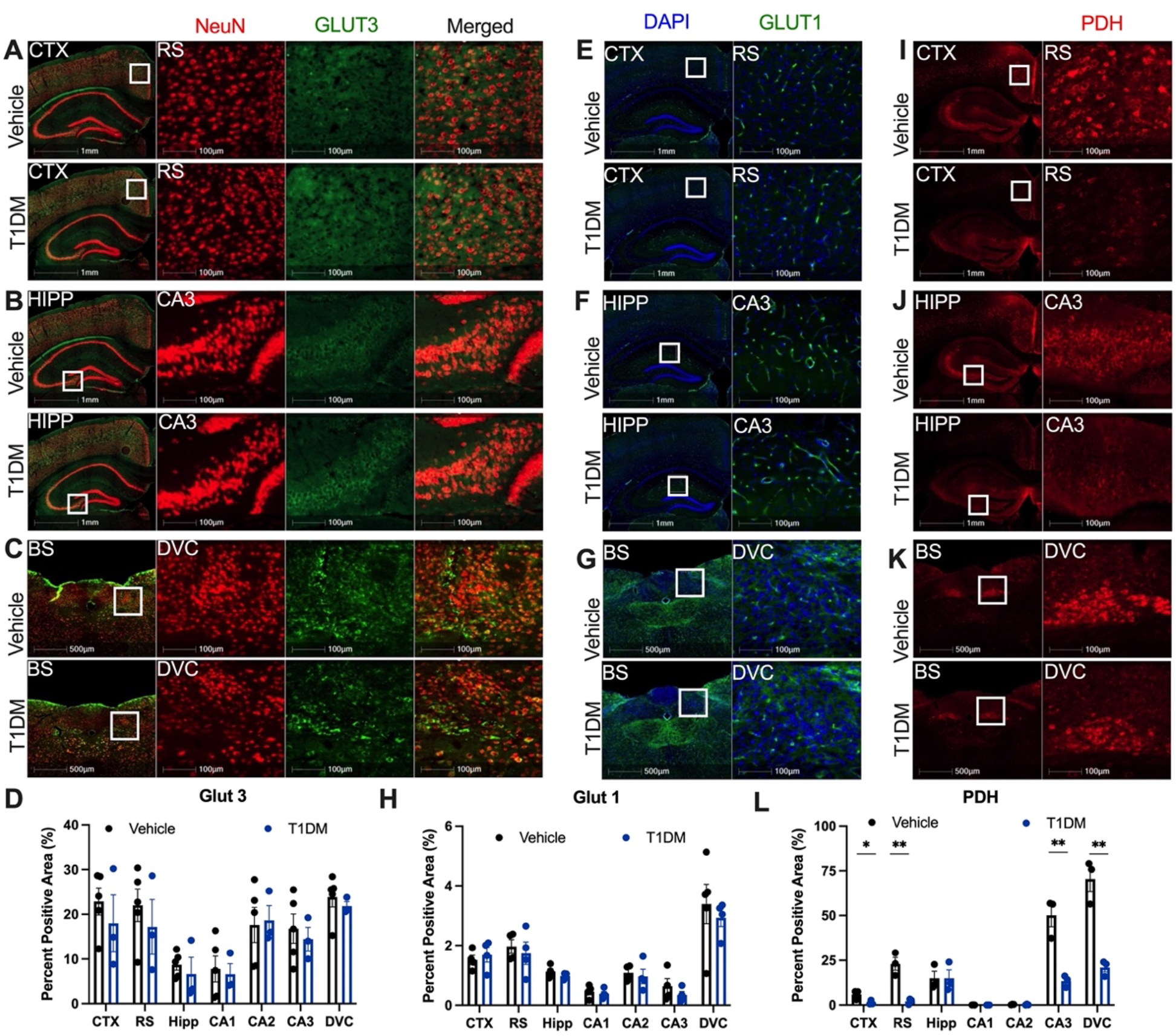
Immunofluorescent analysis of GLUT3, GLUT1 and PDH expression in neocortex (CTX), hippocampus (HIPP) and brainstem (BS) in a mouse model of T1DM. (**A-D**) 20μm coronal sections of the mouse brain were stained with GLUT3 (green) and NeuN (red) and quantified in the CTX (**A**), HIPP (**B**), and BS (**C**). Zoomed-in images are shown for the retro-splenial cortex (RS), Cornu Ammonis 3 (CA3) of HIPP, and dorsal vagal complex (DVC) of BS. (**D**) Quantification of % positive pixels for Glut3 using HALO software, area of interest include whole cortex, RS (zoomed in), whole HIPP, CA1-CA3 (zoomed in), and DVC following mouse brain atlas, and quantification was run for positive pixels in each channel and colocalization. Values are presented as mean ±SEM (n=4 biological replicates), *p<0.05, **p<0.01, ***p<0.001, analyzed by student t-test. (**E-H**) 20μm coronal sections of the mouse brain were stained with GLUT1 (green) and DAPI (blue) and quantified in the CTX (**E**), HIPP (**F**), and BS (**G**). Zoomed-in images are shown for the RS, CA3 of HIPP, and DVC of BS. (**H**) Quantification of % positive pixels for GLUT1 using HALO software, area of interest including whole neocortex, RS (zoomed in), whole HIPP, CA1-CA3 (zoomed in), and DVC following mouse brain atlas, and quantification was run for positive pixels in each channel and colocalization. Values are presented as mean ±SEM (n=4 biological replicates), *p<0.05, **p<0.01, ***p<0.001, analyzed by student t-test. (**I-L**) 20μm coronal sections of the mouse brain were stained with PDH (red) and quantified in the CTX (**I**), HIPP (**J**), and BS (**K**). Zoomed-in images are shown for the retro-splenial cortex (RS), CA3 of HIPP, and DVC of BS. (**L**) Quantification of % positive pixels for PDH using HALO software, area of interest include whole cortex, RS (zoomed in), whole HIPP, CA1-CA3 (zoomed in), and DVC following mouse brain atlas, and quantification was run for positive pixels in each channel and colocalization. Values are presented as mean ±SEM (n=4 biological replicates), *p<0.05, **p<0.01, ***p<0.001, analyzed by student t-test. Scale bars are either 1mm, 500μm, or 100μm. GLUT1-glucose transporter 1, GLUT3-glucose transporter 3, PDH-pyruvate dehydrogenase.

## References

1 Erbsloh, F., Bernsmeier, A. & Hillesheim, H. The glucose consumption of the brain & its dependence on the liver. Arch Psychiatr Nervenkr Z Gesamte Neurol Psychiatr 196, 611–626 (1958).

2 Díaz-García, C. M. & Yellen, G. Neurons rely on glucose rather than astrocytic lactate during stimulation. Journal of neuroscience research 97, 883–889 (2019).

3 Dienel, G. A. Brain glucose metabolism: integration of energetics with function. Physiological reviews 99, 949–1045 (2019).

4 Nimgampalle, M., Chakravarthy, H. & Devanathan, V. in Recent Developments in Applied Microbiology and Biochemistry 77–88 (Elsevier, 2021).

5 Sibson, N. R. et al. In vivo13C NMR measurement of neurotransmitter glutamate cycling, anaplerosis and TCA cycle flux in rat brain during [2-13C] glucose infusion. Journal of neurochemistry 76, 975–989 (2001).

6 Sun, R. C. et al. Noninvasive liquid diet delivery of stable isotopes into mouse models for deep metabolic network tracing. Nature communications 8, 1–10 (2017).

7 Conroy, L. R., Hawkinson, T. R., Young, L. E., Gentry, M. S. & Sun, R. C. Emerging roles of N-linked glycosylation in brain physiology and disorders. Trends in Endocrinology & Metabolism 32, 980–993 (2021).

8 Telser, A., Robinson, H. & Dorfman, A. The biosynthesis of chondroitin sulfate. Archives of Biochemistry and Biophysics 116, 458–465 (1966).

9 Sugahara, K. & Kitagawa, H. Heparin and heparan sulfate biosynthesis. IUBMB life 54, 163–175 (2002).

10 Mosconi, L. et al. Multicenter standardized 18F-FDG PET diagnosis of mild cognitive impairment, Alzheimer’s disease, and other dementias. Journal of nuclear medicine 49, 390–398 (2008).

11 Walker, Z. et al. Clinical utility of FDG PET in Parkinson’s disease and atypical parkinsonism associated with dementia. European Journal of Nuclear Medicine and Molecular Imaging 45, 1534–1545 (2018).

12 Willmann, O., Wennberg, R., May, T., Woermann, F. & Pohlmann-Eden, B. The contribution of 18F-FDG PET in preoperative epilepsy surgery evaluation for patients with temporal lobe epilepsy: a meta-analysis. Seizure 16, 509–520 (2007).

13 Pascual, J. M., Van Heertum, R. L., Wang, D., Engelstad, K. & De Vivo, D. C. Imaging the metabolic footprint of Glut1 deficiency on the brain. Annals of Neurology: Official Journal of the American Neurological Association and the Child Neurology Society 52, 458–464 (2002).

14 Baranwal, A., Mirbolooki, M. R. & Mukherjee, J. Initial Assessment of β3-Adrenoceptor-Activated Brown Adipose Tissue in Streptozotocin-Induced Type 1 Diabetes Rodent Model Using [18F]Fluorodeoxyglucose Positron Emission Tomography/Computed Tomography. Mol Imaging 14, 22–33 (2015).

15 Monsorno, K., Buckinx, A. & Paolicelli, R. C. Microglial metabolic flexibility: emerging roles for lactate. Trends in Endocrinology & Metabolism (2022).

16 Zhao, Y. & Xu, H. Microglial lactate metabolism as a potential therapeutic target for Alzheimer’s disease. Molecular Neurodegeneration 17, 1–3 (2022).

17 Hawkinson, T. R. et al. In Situ Spatial Glycomic Imaging of Mouse and Human Alzheimerżs Disease Brains. Alzheimer’s & Dementia In Press (2021).

18 Jakkamsetti, V. et al. Brain metabolism modulates neuronal excitability in a mouse model of pyruvate dehydrogenase deficiency. Science translational medicine 11, eaan0457 (2019).

19 Muccioli, L. et al. FDG-PET assessment and metabolic patterns in Lafora disease. European Journal of Nuclear Medicine and Molecular Imaging 47, 1576–1584 (2020).

20 Sun, R. C. et al. Brain glycogen serves as a critical glucosamine cache required for protein glycosylation. Cell Metabolism 33, 1404–1417. e1409 (2021).

21 Maher, E. A. et al. Metabolism of [U-13C] glucose in human brain tumors in vivo. NMR in biomedicine 25, 1234–1244 (2012).

22 Choi, C. et al. 2-hydroxyglutarate detection by magnetic resonance spectroscopy in IDH-mutated patients with gliomas. Nature medicine 18, 624–629 (2012).

23 Merritt, M. E. et al. Hyperpolarized 13C allows a direct measure of flux through a single enzyme-catalyzed step by NMR. Proceedings of the National Academy of Sciences 104, 19773–19777 (2007).

24 Ghini, V. et al. Metabolomics profiling of pre-and post-anesthesia plasma samples of colorectal patients obtained via Ficoll separation. Metabolomics 11, 1769–1778 (2015).

25 Lei, H., Duarte, J. M., Mlynarik, V., Python, A. & Gruetter, R. Deep thiopental anesthesia alters steady-state glucose homeostasis but not the neurochemical profile of rat cortex. J Neurosci Res 88, 413–419 (2010).

26 Brewer, M. K. et al. Targeting pathogenic Lafora bodies in Lafora disease using an antibody-enzyme fusion. Cell metabolism 30, 689–705. e686 (2019).

27 Duran, J. et al. Astrocytic glycogen accumulation drives the pathophysiology of neurodegeneration in Lafora disease. Brain 144, 2349–2360 (2021).

28 Friede, R. L. & van Houten, W. H. Relations between post-mortem alterations and glycolytic metabolism in the brain. Experimental Neurology 4, 197–204 (1961).

29 Rauckhorst, A. J. et al. Mouse tissue harvest-induced hypoxia rapidly alters the in vivo metabolome, between-genotype metabolite level differences, and 13C-tracing enrichments. bioRxiv (2022).

30 Suarez, R., Staples, J., Lighton, J. & West, T. Relationships between enzymatic flux capacities and metabolic flux rates: nonequilibrium reactions in muscle glycolysis. Proceedings of the National Academy of Sciences 94, 7065–7069 (1997).

31 Slavov, N., Budnik, B. A., Schwab, D., Airoldi, E. M. & van Oudenaarden, A. Constant growth rate can be supported by decreasing energy flux and increasing aerobic glycolysis. Cell reports 7, 705–714 (2014).

32 Shestov, A. A. et al. Quantitative determinants of aerobic glycolysis identify flux through the enzyme GAPDH as a limiting step. elife 3, e03342 (2014).

33 French, T., Goode, A. & Sugden, M. Ischaemia and tissue pyruvate dehydrogenase activities in the rat: a comparison of the effects of cervical dislocation and pentobarbital anaesthesia. Biochemistry International 13, 843–852 (1986).

34 Zalewska, T. & Domanska-Janik, K. Energy utilization and changes in some intermediates of glucose metabolism in normal and hypoxic rat brain after decapitation. Resuscitation 7, 199–206 (1979).

35 Overmyer, K. A., Thonusin, C., Qi, N. R., Burant, C. F. & Evans, C. R. Impact of anesthesia and euthanasia on metabolomics of mammalian tissues: studies in a C57BL/6J mouse model. PloS one 10, e0117232 (2015).

36 DiNuzzo, M. et al. State-dependent changes in brain glycogen metabolism. Brain Glycogen Metabolism, 269–309 (2019).

37 O’Callaghan, J. P. & Sriram, K. Focused microwave irradiation of the brain preserves in vivo protein phosphorylation: comparison with other methods of sacrifice and analysis of multiple phosphoproteins. Journal of neuroscience methods 135, 159–168 (2004).

38 English, N. J. & MacElroy, J. Molecular dynamics simulations of microwave heating of water. The Journal of chemical physics 118, 1589–1592 (2003).

39 Mayers, C. Histological fixation by microwave heating. Journal of Clinical Pathology 23, 273 (1970).

40 Login, G. Microwave fixation versus formalin fixation of surgical and autopsy tissue. The American Journal of Medical Technology 44, 435–437 (1978).

41 Boychuk, C. R. & Smith, B. N. Glutamatergic drive facilitates synaptic inhibition of dorsal vagal motor neurons after experimentally induced diabetes in mice. J Neurophysiol 116, 1498–1506 (2016).

42 Halmos, K. C. et al. Molecular and functional changes in glucokinase expression in the brainstem dorsal vagal complex in a murine model of type 1 diabetes. Neuroscience 306, 115–122 (2015).

43 Andres, D. A. et al. Improved workflow for mass spectrometry–based metabolomics analysis of the heart. Journal of Biological Chemistry 295, 2676–2686 (2020).

44 Williams, H. C. et al. Oral gavage delivery of stable isotope tracer for in vivo metabolomics. Metabolites 10, 501 (2020).

45 Young, L. E. A. et al. Accurate and sensitive quantitation of glucose and glucose phosphates derived from storage carbohydrates by mass spectrometry. Carbohydr Polym 230, 115651 (2020).

46 Pitra, S. & Smith, B. N. Musings on the wanderer: What’s new in our understanding of vago-vagal reflexes? VI. Central vagal circuits that control glucose metabolism. Am J Physiol Gastrointest Liver Physiol 320, G175–g182 (2021).

47 Kaushik, A. K. & DeBerardinis, R. J. Applications of metabolomics to study cancer metabolism. Biochimica et Biophysica Acta (BBA)-Reviews on Cancer 1870, 2–14 (2018).

48 Johnston, K. et al. Isotope tracing reveals glycolysis and oxidative metabolism in childhood tumors of multiple histologies. Med 2, 395–410. e394 (2021).

49 Jolivalt, C. G., Calcutt, N. A. & Masliah, E. Similar pattern of peripheral neuropathy in mouse models of type 1 diabetes and Alzheimer’s disease. Neuroscience 202, 405–412 (2012).

50 Li, W., Huang, E. & Gao, S. Type 1 diabetes mellitus and cognitive impairments: a systematic review. Journal of Alzheimer’s disease 57, 29–36 (2017).

51 Bolla, A. M. et al. Expression of glucose transporters in duodenal mucosa of patients with type 1 diabetes. Acta Diabetol 57, 1367–1373 (2020).

52 Wasik, A. A. & Lehtonen, S. Glucose Transporters in Diabetic Kidney Disease-Friends or Foes? Front Endocrinol (Lausanne) 9, 155 (2018).

53 Jiang, Y. K., Xin, K. Y., Ge, H. W., Kong, F. J. & Zhao, G. Upregulation Of Renal GLUT2 And SGLT2 Is Involved In High-Fat Diet-Induced Gestational Diabetes In Mice. Diabetes Metab Syndr Obes 12, 2095–2105 (2019).

54 Szablewski, L. Glucose transporters in healthy heart and in cardiac disease. Int J Cardiol 230, 70–75 (2017).

55 Suzanne, M. & Tong, M. Brain metabolic dysfunction at the core of Alzheimer’s disease. Biochemical pharmacology 88, 548–559 (2014).

56 Cunnane, S. et al. Brain fuel metabolism, aging, and Alzheimer’s disease. Nutrition 27, 3-20 (2011).

57 McNair, L. F. et al. Metabolic characterization of acutely isolated hippocampal and cerebral cortical slices using [U-13C] glucose and [1, 2-13C] acetate as substrates. Neurochemical Research 42, 810–826 (2017).

58 Lee, D. C. et al. A lactate-induced response to hypoxia. Cell 161, 595–609 (2015).

59 Ganapathy-Kanniappan, S. & Geschwind, J.-F.H. Tumor glycolysis as a target for cancer therapy: progress and prospects. Molecular cancer 12, 1–11 (2013).

60 Cade, W. T. Diabetes-related microvascular and macrovascular diseases in the physical therapy setting. Phys Ther 88, 1322–1335 (2008).

61 Lacy, M. E. et al. Long-term Glycemic Control and Dementia Risk in Type 1 Diabetes. Diabetes Care 41, 2339–2345 (2018).

62 van Golen, L. W. et al. Cerebral blood flow and glucose metabolism measured with positron emission tomography are decreased in human type 1 diabetes. Diabetes 62, 2898–2904 (2013).

63 Basu, A. et al. Effects of type 2 diabetes on the ability of insulin and glucose to regulate splanchnic and muscle glucose metabolism: evidence for a defect in hepatic glucokinase activity. Diabetes 49, 272–283 (2000).

64 Bischof, M. G. et al. Effects of short-term improvement of insulin treatment and glycemia on hepatic glycogen metabolism in type 1 diabetes. Diabetes 50, 392–398 (2001).

65 Hwang, J. J. et al. Glycemic Variability and Brain Glucose Levels in Type 1 Diabetes. Diabetes 68, 163–171 (2019).

66 Krssak, M. et al. Alterations in postprandial hepatic glycogen metabolism in type 2 diabetes. Diabetes 53, 3048–3056 (2004).

67 Kishore, P. et al. Role of hepatic glycogen breakdown in defective counterregulation of hypoglycemia in intensively treated type 1 diabetes. Diabetes 55, 659–666 (2006).

68 Girard, J. Glucagon, a key factor in the pathophysiology of type 2 diabetes. Biochimie 143, 33–36 (2017).

69 Hatting, M., Tavares, C. D. J., Sharabi, K., Rines, A. K. & Puigserver, P. Insulin regulation of gluconeogenesis. Ann N Y AcadSci 1411, 21–35 (2018).

70 Lee, Y., Wang, M. Y., Du, X. Q., Charron, M. J. & Unger, R. H. Glucagon receptor knockout prevents insulin-deficient type 1 diabetes in mice. Diabetes 60, 391–397 (2011).

71 Petersen, K. F., Price, T. B. & Bergeron, R. Regulation of net hepatic glycogenolysis and gluconeogenesis during exercise: impact of type 1 diabetes. J Clin Endocrinol Metab 89, 4656–4664 (2004).

72 Lee, K. W. et al. Behavioral stress accelerates plaque pathogenesis in the brain of Tg2576 mice via generation of metabolic oxidative stress. Journal of neurochemistry 108, 165–175 (2009).

73 Bonhomme, V. et al. Influence of anesthesia on cerebral blood flow, cerebral metabolic rate, and brain functional connectivity. Current Opinion in Anesthesiology 24, 474–479 (2011).

74 Li, S.-Q. et al. Deficiency of macrophage migration inhibitory factor attenuates tau hyperphosphorylation in mouse models of Alzheimer’s disease. Journal of neuroinflammation 12, 1–11 (2015).

75 Ximerakis, M. et al. Single-cell transcriptomic profiling of the aging mouse brain. Nature neuroscience 22, 1696–1708 (2019).

76 Jiang, S. et al. Combined protein and nucleic acid imaging reveals virus-dependent B cell and macrophage immunosuppression of tissue microenvironments. Immunity (2022).

77 Keren, L. et al. MIBI-TOF: A multiplexed imaging platform relates cellular phenotypes and tissue structure. Science advances 5, eaax5851 (2019).

78 Jung, J. Y. & Oh, M. K. Isotope labeling pattern study of central carbon metabolites using GC/MS. J Chromatogr B Analyt Technol Biomed Life Sci 974, 101–108 (2015).

79 MacRae, J. I. et al. Mitochondrial metabolism of glucose and glutamine is required for intracellular growth of Toxoplasma gondii. Cell Host Microbe 12, 682–692 (2012).

80 Fiehn, O., Kopka, J., Trethewey, R. N. & Willmitzer, L. Identification of uncommon plant metabolites based on calculation of elemental compositions using gas chromatography and quadrupole mass spectrometry. Anal Chem 72, 3573–3580 (2000).

81 Kind, T. et al. FiehnLib: mass spectral and retention index libraries for metabolomics based on quadrupole and time-of-flight gas chromatography/mass spectrometry. Anal Chem 81, 10038–10048 (2009).

82 Fiehn, O. Metabolomics by Gas Chromatography-Mass Spectrometry: Combined Targeted and Untargeted Profiling. Curr Protoc Mol Biol 114, 30.34.31–30.34.32 (2016).

83 Dagley, M. J. & McConville, M. J. DExSI: a new tool for the rapid quantitation of 13C-labelled metabolites detected by GC-MS. Bioinformatics 34, 1957–1958 (2018).

84 Heinrich, P. et al. Correcting for natural isotope abundance and tracer impurity in MS-, MS/MS-and high-resolution-multiple-tracer-data from stable isotope labeling experiments with IsoCorrectoR. Sci Rep 8, 17910 (2018).

